# Assessment of neutralization susceptibility of Omicron subvariants XBB.1.5 and BQ.1.1 against broad-spectrum neutralizing antibodies through epitopes mapping

**DOI:** 10.1101/2023.03.01.530717

**Authors:** Masaud Shah, Hyun Goo Woo

**Author notes:** **Corresponding author** Hyun Goo Woo, M.D., Ph.D.

## Abstract

The emergence of new variants of the SARS-CoV-2 virus has posed a significant challenge in developing broadly neutralizing antibodies (nAbs) with guaranteed therapeutic potential. Some nAbs, such as Sotrovimab, have exhibited varying levels of efficacy against different variants, while others, such as Bebtelovimab and Bamlanivimab-etesevimab are ineffective against specific variants, including BQ.1.1 and XBB. This highlights the urgent need for developing broadly active mAbs providing prophylactic and therapeutic benefits to high-risk patients, especially in the face of the risk of reinfection from new variants. Here, we aimed to investigate the feasibility of redirecting existing mAbs against new variants of SARS-CoV-2, as well as to understand how BQ.1.1 and XBB.1.5 can evade broadly neutralizing mAbs. By mapping epitopes and escape sites, we discovered that the new variants evade multiple mAbs, including FDA-approved Bebtelovimab, which showed resilience against other Omicron variants. Our approach, which included simulations, free energy perturbations, and shape complementarity analysis, revealed the possibility of identifying mAbs that are effective against both BQ.1.1 and XBB.1.5. We identified two broad-spectrum mAbs, R200-1F9 and R207-2F11, as potential candidates with increased binding affinity to XBB.1.5 and BQ.1.1 compared to the wild-type virus. Additionally, we propose that these mAbs do not interfere with ACE2 and bind to conserved epitopes on the RBD that are not-overlapping, potentially providing a solution to neutralize these new variants either independently or as part of a combination (cocktail) treatment.

## Introduction

SARS-CoV-2 neutralizing antibodies have thus far played a crucial role in preventing and treating COVID-19, but they can be hindered by viral evolution and the virus’s ability to evade the host immune response [1, 2]. This was particularly demonstrated by the emergence of highly contagious BA.1 sublineage in November 2021 and several other variants of concern (VOCs) since the start of the pandemic [3]. The evolution of the Omicron BA.2, BA.4, and BA.5 lineages has led to the emergence of new subvariants, including BA.2.75.2, BA.4.6, BQ.1.1, and recent XBB.1.5 in the US [4], which are highly transmissible and can evade the immune system even in vaccinated individuals [3, 5, 6]. Approximately 80% of the population has been infected with at least one of the Omicron subvariants within a year, due to the lack of effective vaccination [3, 7, 8]. Recent studies have shown that the Omicron subvariants are escaping from neutralization induced by current vaccines, raising concerns about their potential to infect individuals who have received three or four vaccine doses, including a bivalent booster [1, 7, 8]. The new subvariants, particularly BQ.1.1 and XBB.1.5 are expected to become prevalent in many countries by early 2023 due to their additional mutations in the spike.

To be ready for future variants and sarbecovirus pandemics, it is necessary to develop broad-spectrum antibody therapeutics and vaccines. However, we still lack a complete understanding of the Spike epitopes that can induce broad sarbecovirus neutralization. In response to the escalation of the COVID-19 pandemic, many initiatives have been launched to find treatments, including studies on existing medications. Sharing information and resources will help explore potential solutions and increase the chances of finding an immediate and lasting treatment.

A recent cohort study of 936 COVID-19 convalescent patients identified a subset of individuals as “elite neutralizers” with broad-spectrum neutralizing antibodies (broad-nAbs) that could neutralize SARS-CoV-2 VOCs and Omicron subvariants including BA.5 [9]. While some of these monoclonal antibodies could neutralize the subvariants, others escaped due to single-point mutations in the spike [10]. Using our expertise in computational antibody design, we have created models of the broad-nAbs and mapped their conserved epitopes on the RBD. This comprehensive mapping of conserved sites provides important guidelines for the development of broad-spectrum therapeutics against BQ.1.1, XBB.1.5, and other emerging variants.

## Results

### RBD class designation of the broad-nAbs

The RBD-binding antibodies are structurally characterized into 4 classes based on their binding epitopes, their ability to bind an ‘‘up’’ or ‘‘down’’ RBD conformation, and interference with ACE2 binding [9]. Class I antibodies such as C102 block ACE2, bind only to the “up” RBD conformation, and have relatively shorter CDRH3 loops [11]. While, Class II antibodies bind to both “up” and “down” RBD conformations, interact with adjacent RBDs, and neutralize the Spike-ACE2 interaction (**Figure 1A**). Class III antibodies bind to the outside of the ACE2 site, while Class IV antibodies do not block ACE2 and bind only to the “up” RBD conformation [11]. It has been shown that Class I antibodies with short CDRH3s and class II with long CDRH3s are typically knocked out by Lys417 or Glu484 mutants, respectively [12, 13]. However, the current cohort study has identified several IGHV3-53 antibodies that defy this proposed paradigm [14]. These antibodies, with 93.5%–97.3% germline identity, have demonstrated resistance to the typical variant escape due to minor differences in their antibody sequences [9].

**Figure 1.**
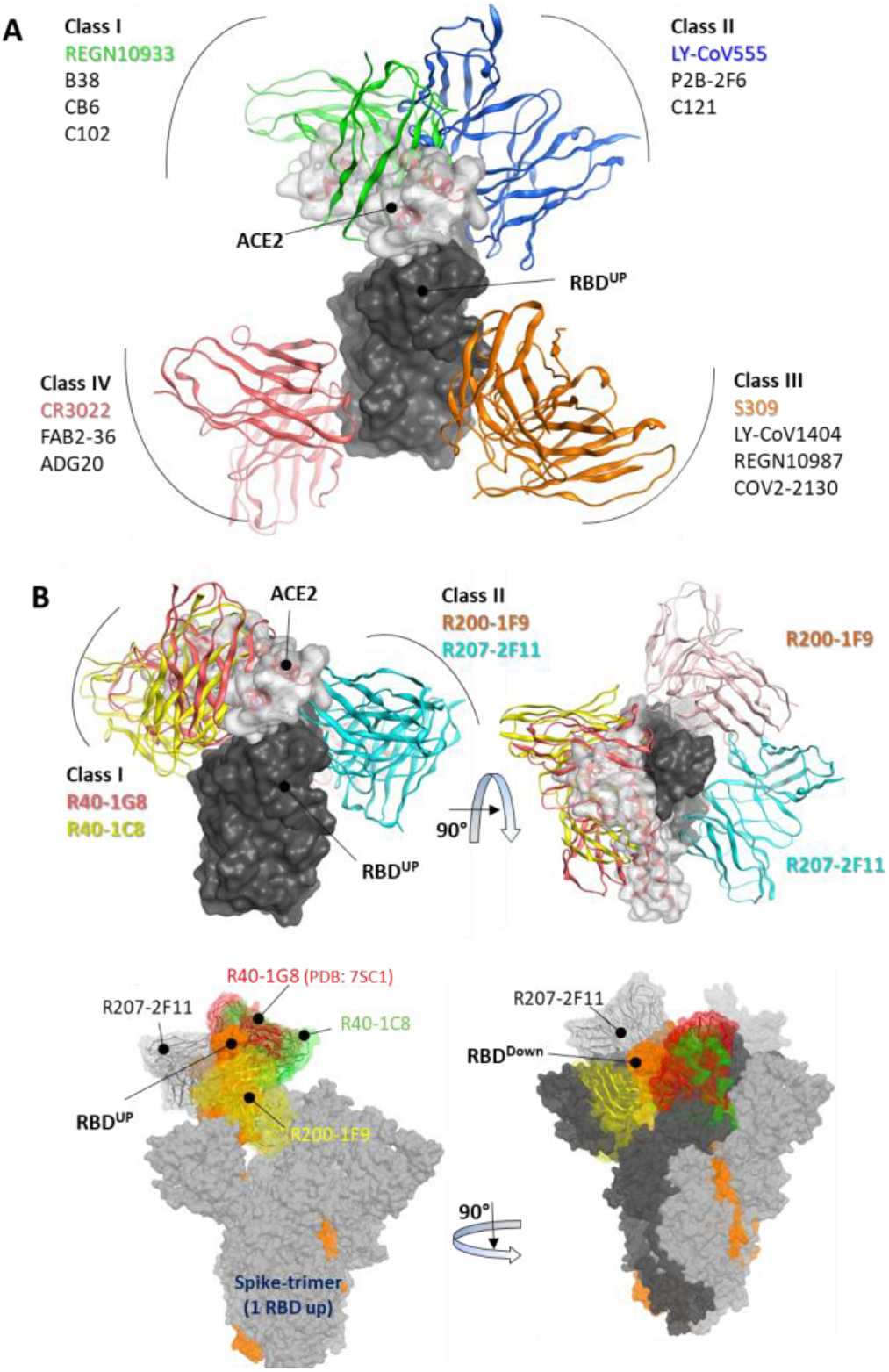
Epitope-based Classification of RBD-Binding Antibodies and Class Designation of Broad-Neutralizing Antibodies (nAbs). (A) Class I and Class II Monoclonal Antibodies (mAbs) interfere with the Angiotensin-Converting Enzyme 2 (ACE2) receptor (represented by the transparent surface), while Class III and Class IV mAbs bind outside the ACE2 interface. (B, *top*) The mAbs R40-1G8 and R40-1C8 directly compete with the ACE2 receptor, while R200-1F9 and R207-2F11 do not. (B, *bottom*) All four mAbs are bound to the trimeric Spike (wild type) protein. Only R207-2F11 accommodates both the up and down conformations, while the other mAbs bind exclusively to the RBD down conformation.

Three broad-nAbs R40-1G8, R40-1C8, and R207-2F11 investigated here are encoded by IGHV3-53 but different light V genes (KV1-9, KV1-9, and KV1-33, respectively). Only R200-1F9 is encoded by the IGHV3-48 gene with longer CDRH3 (**Table 1**). Although R40-1G8 and R207-2F11 share identical CDRH3 and logically they should follow the typical class I rule, both antibodies were found to respond differently to SARS-CoV-2 variants in terms of neutralization. R40-1G8 failed to neutralize all previous Omicron variants while R207-2F11 retained its neutralization capacity [9]. In addition, our epitope mapping suggests that R40-1C8 overlaps with R40-1G8 and partially competes with ACE2 (**Figure 1B**), yet the former neutralized all but BA.4/5 variant of Omicron while the latter was devoid of this potential [14]. This notion suggests that while many IGHV3-53 encoded antibodies may agree to the epitope-based classification of RBD-binding mAbs, some defy this rule.

**Table 1:**
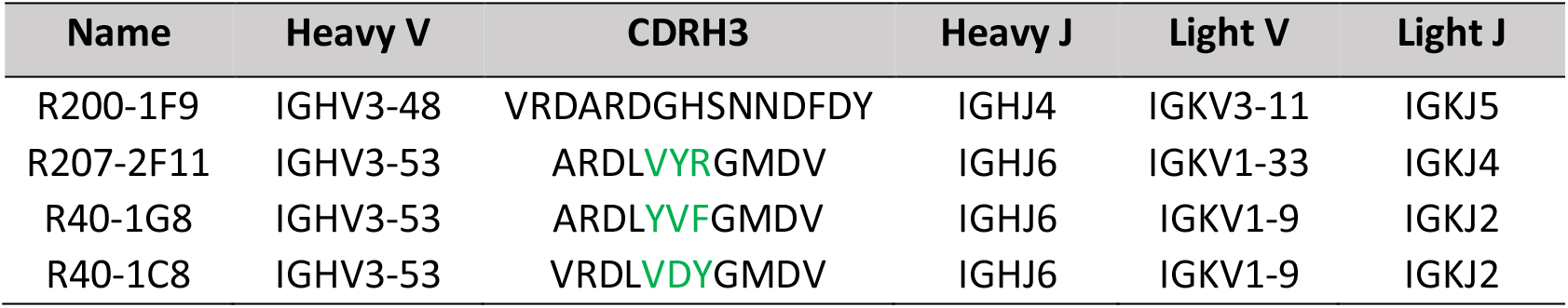
SARS-CoV-2 broad-nAbs and their heavy and light chains encoding genotypes.

After extensive Ag-Ab docking simulations and epitope mapping (discussed below), we designated R40-1G8 and R40-1C8 as class I and R200-1F9 and R207-2F11 as class II antibodies. Both R40-1G8 and R40-1C8 compete with ACE2 while R200-1F9 and R207-2F11 do not (**Figure 1B**). Subsequently constructing full-length trimeric Spike-mAbs models, we propose that R207-2F11 with shorter CDRH3 can bind RBD in both up and down conformation without making any clash with the adjacent RBD, while R200-1F9 with longer CDRH3 may bind the up conformations only (**Figure 1B**). In their cryo-EM analysis, Vanshylla et al. proposed that R40-1G8 Fab bind to both up (state 1) and down (state 2) RBDs; however, due to the relatively low resolution for the RBD and Fab in state 2, they could build a model for state 1 only [9]. Based on our trimeric Spike models and the reported epitope of R40-1G8, we demonstrated that it is not plausible for this mAb to bind the down conformation of RBD, regardless of the up or down conformation of the nearby RBD (discussed in detail below). Based on their non-overlapping epitopes on RBD and the fact that R207-2F11 can potentially bind to both up and down conformation, R40-1C8, R200-1F9, and R207-2F11 could be used as cocktail therapy as they hold potential neutralization against Omicron and other variants of SARS-CoV-2 [14].

### All Omicron subvariants escape R40-1G8 broad-nAb

R40-1G8, one of the broad-nAbs and classified as class I can potentially neutralize Wu01, several other SARS-CoV-2 variants, and 17 variants with single amino acid mutations at the potential RBD escape sites [9]. Interestingly, most of these mutations are now reported in the emerging Omicron variants (**Figure 2A**). However, Florian Klein’s group has recently reported in their pre-print study that all Omicron variants are showing some resistance to R40-1G8. This suggests that R40-1G8 can potentially endure the interface changes brought by a single mutation; however, multiple mutations in the same epitope may force conformation changes in the RBD that obscure the Ag-Ab interface and CDRs-fitting onto their epitopes.

**Figure 2.**
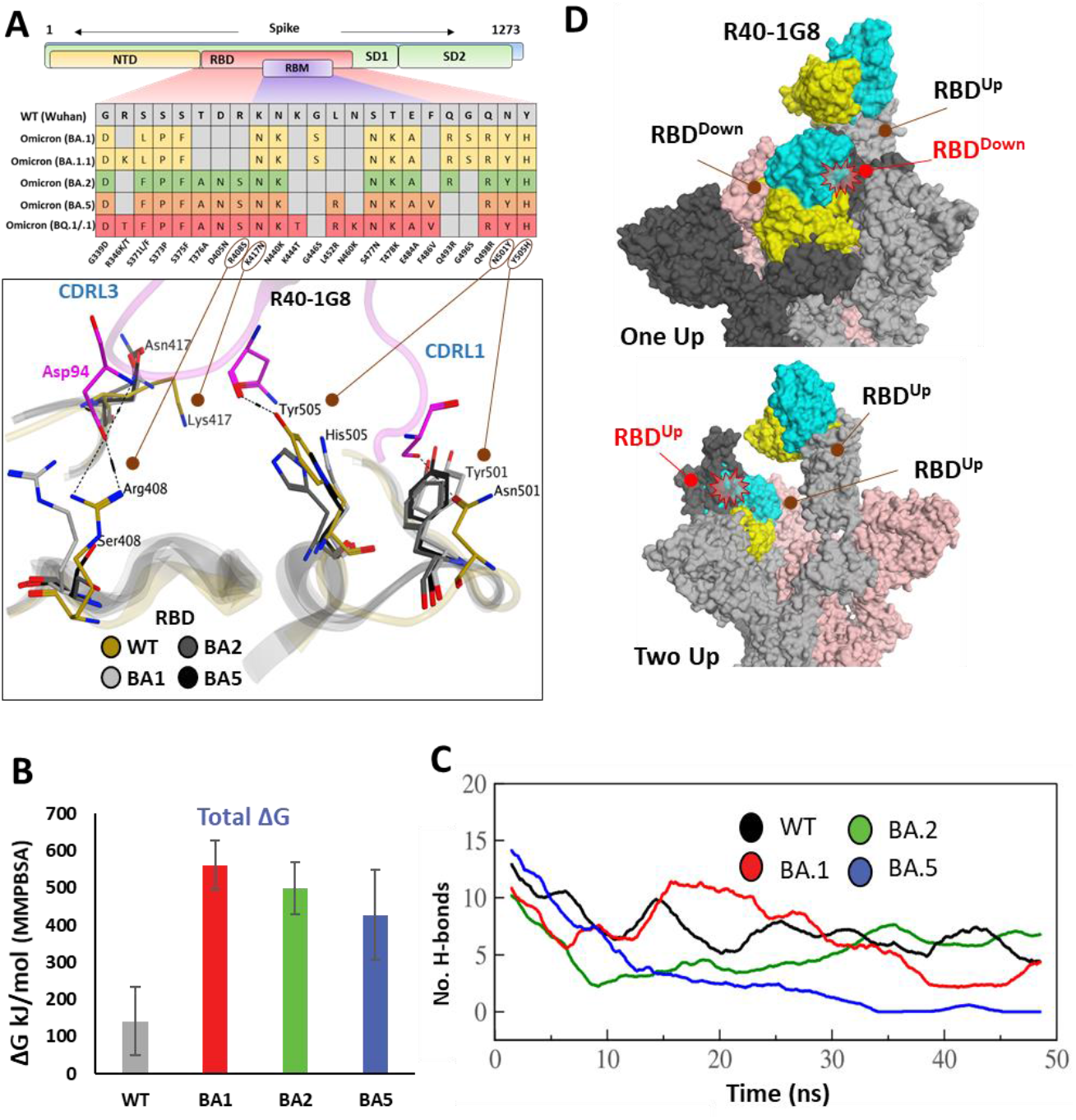
The escape of Omicron subvariants from broad-nAb R40-1G8. (A) Mutations in the RBD region of Spike protein of SARS-CoV-2 Omicron variants and their effect on the interface contacts with respect to R40-1G8. Multiple mutations at the interface are indicated with stick and circles (B) Changes in the binding free energy (BFE) of R40-1G8 bound to the SARS-CoV-2 variants. (C) Changes in the number of Hydrogen bonds between R40-1G8 and respective variants. (D) The R40-1G8 Antibody can only be accommodated onto the RBD in its “up” conformation in the trimeric Spike model. The Fab bound to the down conformation of the RBD clashes with the adjacent RBD in both the up and down states.

To expand upon the binding mode and investigate the escape of Omicron subvariants from this broad-nAb, we constructed their protein models and explored their interfaces. As discussed above, R40-1G8 can resist single mutation at Lys417 and other positions; however, a single amino acid variant at Arg408 was not investigated which appeared in the BA.2, BA.4/5, and BQ.1/.1 variants as Arg408Ser (**Figure S1A**). This is the only amino acid that established an electrostatic bond with the CDRL2 Asp94 and apprehended the R40-1G8-RBD complex, which is otherwise lost in Lys417/Asn and Arg408/Ser double mutant in BA.2, BA.4/5, BQ.1/.1, and XBB.1.5 variants (**Figure 2A** & **Table S1**). The idea that Lys417/Asn single variant did not show resistance to R40-1G8 in the previous findings [9] was due to the contribution of the strong electrostatic bond by Arg408 and therefore defied the typical class I antibodies knockout phenomena by Lys417 mutation [11, 15].

We took advantage of the molecular dynamics simulation (MDS) and binding free energy (BFE) perturbation to validate this loss in the neutralization potential of R40-1G8 against Omicron and its subvariants. The average total binding free energy of 200 structural representatives sampled from a 50ns trajectory suggests that R40-1G8-RBD complexes of the BA.1, BA.2, and BA.5 subvariants of Omicron gain considerable energy along the course of simulation (**Figure 2B**). This increase in the total BFEs indicates the destabilization of the Ag-Ab complexes, which could be attributed to the rise in electrostatic energy (ELE); in other words, the rise in ELE is due to the omission of crucial Arg408-Asp94 salt bridge upon Arg408Ser mutation (**Figure S1B**). Although MMPBSA-based BFE estimation is quite robust [16], we went on to confirm this change in energy by single frame MMGBSA method [17]. The total BFE of RBDWT-nAb was recorded as -110.03 kcal/mol and that of BA.1, BA.2 and BA.5 were -71.97 kcal/mol, -93.66 kcal/mol, and -63.37 kcal/mol, respectively. Only RBDWT-nAb had ELE as -99.52 kcal/mol while other complexes had this energy term over 157.03 kcal/mol. This was further supported by the sudden decline in the hydrogen bonds between nAb and BA.2 and BA.5 RBDs containing Arg408Ser substitution **Figure 2C**). To track this complete loss in the hydrogen bonds network between RBD^BA.5^-nAb, we sampled 1000 frames from the 50ns MDS trajectory and looked for the RBD-Ab separation as a function of time. The fab and RBD molecules completely separated in RBD^BA.5^-nAb but remained intact in RBD^WT^-nAb (GIF S1).

Generally, Class I RBD antibodies can bind to only up conformation of the RBD [18, 19]; surprisingly, R40-1G8 has been reported to bind both up and down conformation of RBD in a trimeric Spike model. In their explanation, Vanshylla et al. suggest that “in the R40-1G8-spike map, there is one ‘‘up’’ RBD with R40-1G8-Fab, one ‘‘up’’ RBD without R40-1G8-Fab, and a ‘‘down’’ RBD with R40-1G8-Fab” [9]. To debate this further, we constructed two trimeric Spike models with two R40-1G8 Fabs; (1) containing one RBD^Up^ bound to the Fab and one of the two RBD^Down^ bound to the second Fab, (2) containing two RBD^Up^ with one empty and the other bound to Fab, and the third RBD^Down^ was bound to Fab (**Figure 2D**). We could demonstrate that R40-1G8 Fab feely binds to the RBD^Up^ conformation; however, following the epitope suggested by Vanshylla et al. the second Fab cannot be accommodated and fit onto the RBD^Down^, irrespective of the down or up conformation of the adjacent RBD. In both scenarios, Fab bound to the down RBD clashes with the adjacent RBD, confirming the general concept of class I RBD antibodies (**Figure 2D**). Since these models are based on the actual epitopes suggested by the same groups and therefore dependable, we believe the alternative binding pose of R40-1G8 onto the down RBD could be an artifact or a transient low-affinity epitope on RBD which was captured during cryo-EM scanning. Overall, these data suggest that all Omicron variants escape from an R40-1G8 due to the collective contribution of multiple mutations in the conformational rearrangement of a relatively conserved epitope.

### R40-1C8 neutralize all SARS-CoV-2 variants except BA.4/5 BQ.1.1 and XBB.1.5

As discussed in the class designation, R40-1C8 is a class I RBD nAb and potentially binds to its up conformation only, partially competing with R40-1G8 but not Bebtelovimab (an FDA-approved COVID-19 broad-spectrum therapeutic mAb) (**Figure 3A**). However, neutralization assay suggests that, unlike R40-1G8, R40-1C8 exhibits significant neutralization against all Omicron sub-variants (IC50 <0.085 μM/mol) but BA.4/5 [9, 14]. Three positively charged residues Arg408, Lys417, and Asp420 can potentially establish salt bridges with the CDRH3; here only the latter two could make such bonds (**Figure 3B** & **Table S2**). The loss of Lys417 salt bridge by BA.1 and BA.2 but their susceptibility to R40-1C8 neutralization suggest that Asp420-CDRH3 contact compensates for this loss and holds the Ag-Ab complex intact. A somewhat similar effect was previously reported, where R40-1G8 was not affected by the K417E/N/T mutation, which is a prominent escape site found in VoCs like B.1.351 [9].

**Figure 3:**
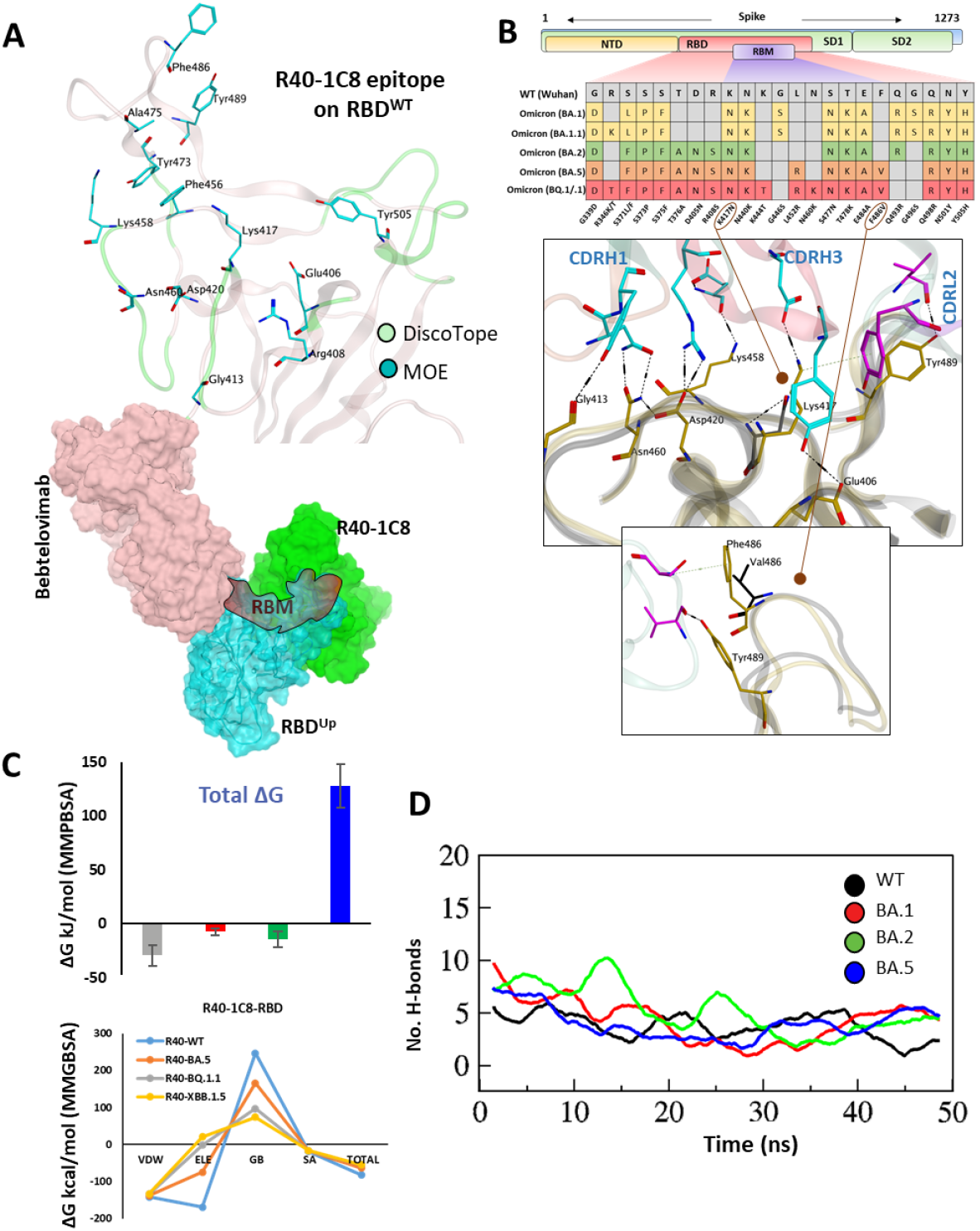
The epitope mapping and escape of BA.5, BQ.1.1, and XBB.1.5 subvariants from R40-1C8. (A) The epitope residues on RBD are predicted through DiscoTope (green color cartoon representation) and Molecular Operating Environment (MOE, Cyan color). The surface maps (colored according to chains) shows that R40-1C8 does not compete with Bebtelovimab. B) Mutations in the RBD region of Spike protein of SARS-CoV-2 Omicron variants and their effect on the interface contacts with respect to R40-1C8. Mutations (including Phe(F)486Val(V)) at the interface are indicated with stick and circles. (C) Changes in the binding free energy (BFE, *top* MMPBSA, *bottom* MMGBSA) of R40-1C8 bound to the SARS-CoV-2 variants. (D) Changes in the number of Hydrogen bonds between R40-1G8 and respective variants.

Considering the current epitope into account, BA.2 and BA.4/5 differ by one residue Phe486Val situated in the far flexible loop of the RBM motif of RBD, which establishes an auxiliary π-cation bond with Ser60 near CDRL2 (**Figure 3A, B**), yet BA.2 but not BA.5 is neutralized by R40-1C8. Our previous findings suggest that this loop in RBM is highly flexible and plays a crucial role in ACE2 recognition and binding [20]. The potential role of Phe486Val mutation and its resistance to R40-1C8 neutralization was further validated by the huge difference in total BFE of BA.5 against other Omicron sub-variants and RBD^WT^ as well (**Figure 3C**). Further investigation suggested that this rise in total BFE (in other words destabilization of the R40-1C8-RBD^BA.5^ complex) could be attributed to the loss in ELE potential (**Figure S1C**). This was further validated by MMGBSA-based BFE calculation where the ELE potential raised from -169.33 kcal/mol (RBDWT-R40-1C8) to -74.77 kcal/mol (RBD^BA.5^-R40-1C8). We went on to investigate the interface alteration induced by BQ.1.1 and XBB.1.5 variants and calculated the loss in binding affinity. Like BA.5 where the total binding affinity dropped by ∼21.0 kcal/mol, the affinity of both BQ.1.1 and XBB.1.5 towards R40-1C8 dropped by 28 kcal/mol (**Figure 3C**, *bottom*). Surprisingly, the electrostatic potentials in both cases dropped by over 170 kcal/mol. This change in affinity was calculated by the MMGBSA method.

To look further, we extracted 1000 frames from the 50ns MDS trajectory and considered the relative positioning of Phe486 in BA.1 and Val486. The heavy chain of antibody remained intact in BA.1-R40-1C8 complex but dissociated in BA.5-R40-1C8 complex (GIF S2), suggesting that in addition to electrostatic contacts, Van der Waals forces also play a crucial role in Ag-Ab stability. Upon hydrogen bond network investigation, there was a steep decline in overall hydrogen bonds till 30ns between BA.5-R40-1C8 complexes, which was slightly restored in the last quarter of the MDS course, perhaps due to new bonds formed between the VL chain and RBD (**Figure 1D**). Altogether, the epitope mapping and BFE perturbation suggest that Omicron sub-variants like BA.5, BQ.1/.1, and XBB.1.5 containing Phe->Val/Pro mutations are highly likely to escape the R40-1C8 neutralization.

### R200-1F9 and R207-2F11 neutralize all Omicron variants

As discussed above, R200-1F9 can potentially bind to the RBD^Up^ only without competing with ACE2, whereas R207-2F11, a class II RBD nAb can potentially bind a conserved epitope on RBD in both ‘up’ and ‘down’ conformation. A co-model suggests that both R200-1F9 and R207-2F11 could be used as cocktail therapy with Bebtelovimab as they do not compete on RBD (Figure 4A). Since both R207-2F11 and R200-1F9 do not interfere with ACE2 directly, they possibly neutralize the virus by restricting the flexibility of RBM as most of their epitope residues are located adjacent to this motif (**Figure S2**). Interface analyses indicate that the R200-1F9-RBD complex is stabilized by two salt bridges between Arg466 and Glu471 and CDHR2, which remains unchanged in all Omicron sub-variants (**Table S3**). However, His104 in CDRH3 creates a hydrogen bond with the backbone nitrogen of Asn460 which has been reported to be mutated into Lys460 in BQ.1.1 and XBB.1.5 variants. To explore whether Asn460/Lys affect this interaction, we constructed the RBD models of BQ.1.1 and XBB.1.5 and observed that the backbone nitrogen retains this bond in both cases (**Figure 4B & Table S3)**.

**Figure 4:**
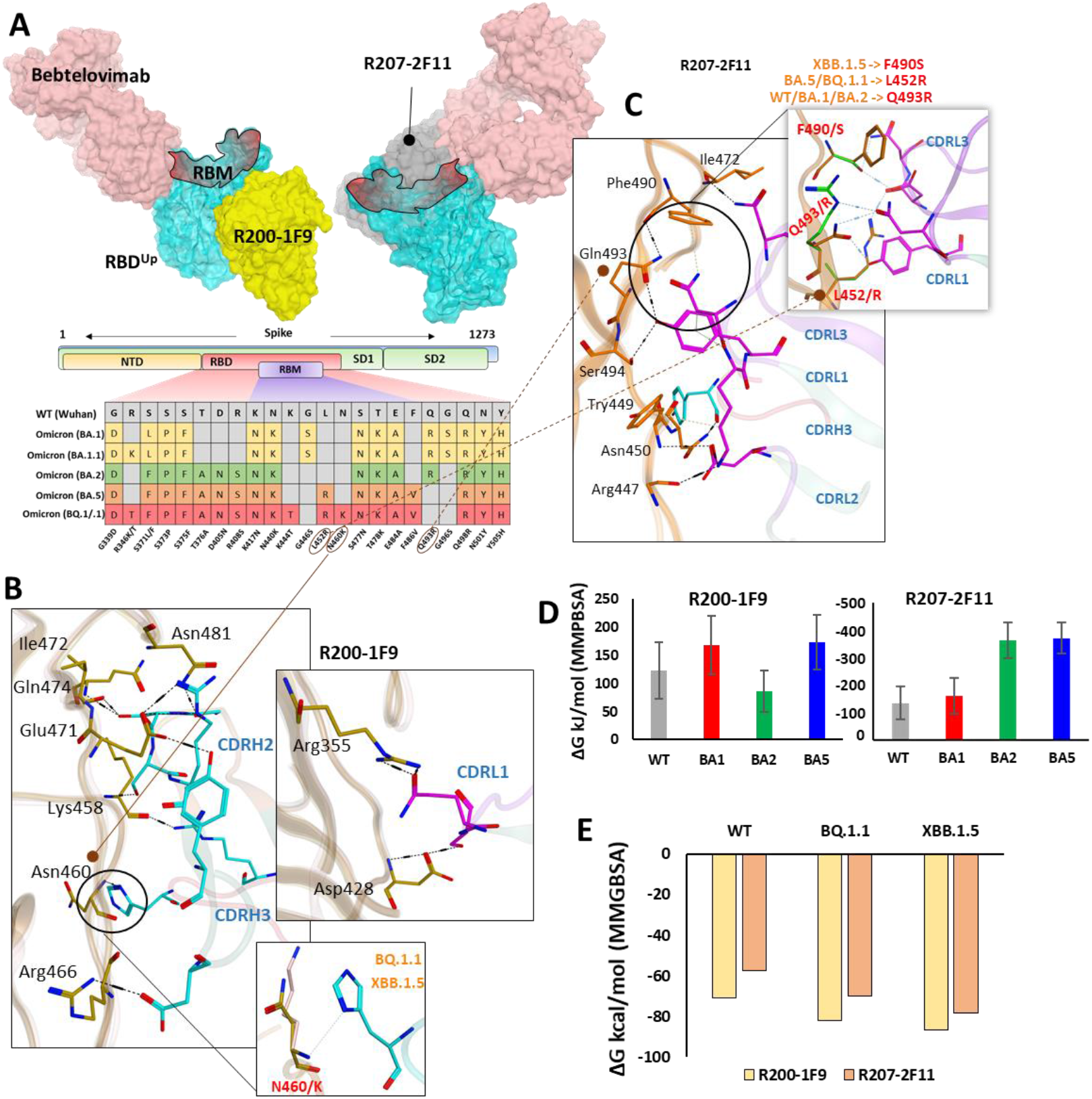
R200-1F9 and R207-2F11 neutralizes all Omicron variant. (A) The surface maps (colored according to chains) shows that both R200-1F9 and R207-2F11 do not compete with Bebtelovimab. Mutations in the RBD region of Spike protein of SARS-CoV-2 Omicron variants and their effect on the interface contacts with respect to R200-1F9 and R207-2F11. (B) Interface residues of the R200-1F9 with respect to Omicron variants. (C) Interface residues of the R207-2F11 with respect to Omicron variants. (D) Changes in the binding free energy (BFE, calculated through MMPBSA) of both R200-1F9 and R207-2F11 bound to the SARS-CoV-2 variants BA.1-B.5. (E) Changes in the binding free energy (BFE, calculated through MMGBSA) of both R200-1F9 and R207-2F11 bound to the SARS-CoV-2 variants BQ.1.1 and XBB.1.5.

Unlike R200-1F9 where two strong electrostatic contacts were involved, most of the interface contacts were hydrogen bonds and other auxiliary forces in the R207-2F11-RBD complex, contributed by all three CDRs in light chains and a single hydrogen bond by CDRH3 (**Figure 4C**). The reason R207-2F11 retained its neutralization effect against all previously reported Omicron sub-variants seems to be due to Arg493 in BA.1 and BA.2 which rather enhanced the Ab-binding energy compared to Gln493 in WT, BA.5, and BQ.1.1 (**Table S4**). This could be one of the driving forces that almost doubled the binding affinity of R207-2F11 against RBD^BA.2^ (Figure 4D). One of the most prominent mutations we observed was Leu452Arg in BA.5 and BQ.1.1 which established a salt bridge with the backbone oxygen of CDRL3 Asp92 (**Table S4**). In addition, the loss of cation-π interaction between CDRL1 Asn30 and RBD Phe490 was substituted by a considerably strong Ser^490^-Asp^92^ hydrogen bond in RBD^XBB.1.5^ (**Figure 4C**). As BA.5 and BQ.1.1 variants contain Leu452Arg and other substitutions that enhance the antibody affinity, we suggest the R207-2F11 broad-nAb potentially retains its neutralization against XBB.1.5 (Leu452 is not mutated) and BQ.1.1 variants; however, this notion may require further experimental validation.

To further investigate the impact of Omicron variants on the binding affinity of these mAbs, we calculated their BFEs. Unlike R40-1C8, which showed a significant decrease in binding affinity for all Omicron variants, and R401G8, which lost affinity only against BA.5, both R200-1F9 and R207-2F11 were found to have a similar or improved binding affinity for all Omicron variants of RBD compared to the RBD^WT^ (**Figure 4D** & **Figure S3**). This data was further validated by MMGBSA-based BFE calculation, where both XBB.1.5 and BQ.1.1 substantially enhanced their overall binding affinity against R207-2F11 and R200-1F9 (**Figure 4E**). Previous studies have shown that most broad-spectrum nAbs remain effective against single amino acid variants in the virus, but multiple mutations in the same epitope can reduce their efficacy. In the cases of the new BQ.1.1 and XBB.1.5 variants, many of the mutations occur outside the binding epitopes of these mAbs. The preservations of BFEs and the nature auxiliary contacts made by single amino acid mutations within the epitopes suggest that these broad-spectrum nAbs may still be effective against the new Omicron variants of BQ.1.1 and XBB.1.5.

### The escape of Omicron BQ.1.1 and XBB.1.5 from Bebtelovimab

Among all available mAbs, Bebtelovimab stands out to be the best and has shown remarkable activity against all SARS-CoV-2 variants that have been reported until recently, including BA.4/5 [21]. However, the emergence of a sub-variant of the BA.5 variant, BQ.1.1, and a sub-variant of the BA.2 variant, XBB, and XBB.1.5 has shown some extraordinary spread across multiple countries including the US and India [5]. Their ancestors have already shown less sensitivity to a broad range of FDA-approved antibodies [22] and additional mutations in the RBD have put the neutralization potential of active nAbs like Bebtelovimab at further risk (Figure 5A).

**Figure 5:**
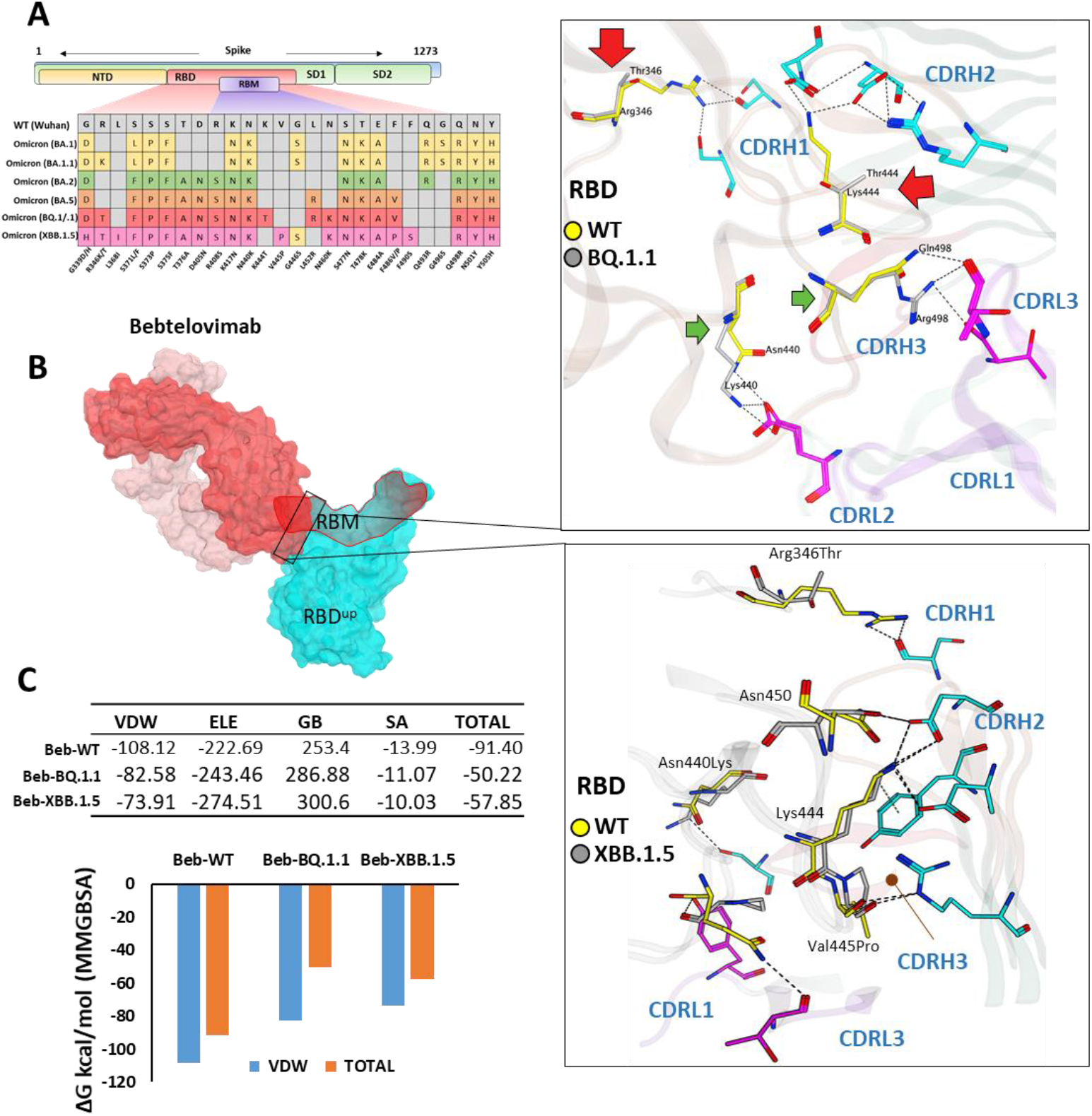
The escape of Omicron BQ.1.1 and XBB.1.5 from Bebtelovimab. (A) Mutations in the RBD region of Spike protein of SARS-CoV-2 Omicron variants and their effect on the interface contacts with respect to Bebtelovimab. (B) Mutations in both BQ.1.1 and XBB.1.5 abolish the Bebtelovimab interface. (C) Changes in the binding free energy (BFE, calculated through MMGBSA). Bebtelovimab loses its binding affinity against both new variants.

To answer this question and predict their potential neutralization escape, we built the RBD models of both BQ.1.1 and XBB.1.5 bound to Bebtelovimab and calculated their BFE and interface changes. The potential of Bebtelovimab to retain its neutralization against broad-range SARS-CoV-2 variants till now is attributed to its relatively conserved epitope on RBD as well as the utilization of all six CDRs in antigen binding (**Figure 5B** & **Table 2**). However, the emergence of BQ.1.1 with multiple mutations within its epitope could render this antibody ineffective. Four substitutions including Arg346Thr, Lys444Thr which abolished the strong electrostatic bonds, and Gln498Arg and Asn440Lys which established relatively stronger contacts with Bebtelovimab are of particular concern. Arg346Thr, Lys444Thr abolishes a network of salt bridges and hydrogen bonds with the CDRH1 and CDRH2 (Figure 5B). Similarly, XBB.1.5 which emerged with multiple new substitutions within the Bebtelovimab epitope, seems to weaken the Ag-Ab interface. Since XBB.1.5 is a descendent of BA.2, the Salt bridges established by Lys444 in RBD^WT^ with CDRH2 remained intact; however, there are two more substitutions Val445Pro and Gly446Ser in the same loop containing Lys444, which may restrict the loop flexibility, resulting in the reduced CDR-fitting. This model, RBD^XBB.1.5^-Bebtelovimab, was subjected to a 100ns MDS to see the interface changes. As suggested, the interface contacts dropped by half (total of 7 hydrogen bonds) as compared to the RBDWT (13 hydrogen bonds) (Table 2). Surprisingly, the MMGBSA-based BFE dropped by 45% in RBD^BQ.1.1^-Bebtelovimab, as compared to RBD^WT^, suggesting that Bebtelovimab escaped this variant. In the case of RBD^XBB.1.5^ the total BFE dropped to ∼36.26% and that of Van der Waal energy dropped by ∼30% (**Figure 5C**). While this study was ongoing, multiple preliminary reports confirmed that both BQ.1.1 and XBB.1.5 escaped many anti-COVID-19 antibodies that are either approved by FDA e.g. Bebtelovimab, or currently undergoing clinical investigations [21, 23]. Here we noticed that, although mutations such as Gln498Arg and Asn440Lys which rather strengthen the Bebtelovimab interface with these variants are surpassed by other mutations like Arg346Thr, Lys444Thr, Val445Pro, and Gly446Ser in one way or another, ultimately abrogating their interfaces, as suggested by the single amino acid energy contribution (**Figure S4**). This further strengthens the concept that single mutations in a relatively conserved epitope could be endured by a broad-nAb; however, multiple mutations induce considerable conformational alterations in the epitopes that are beyond the CDRs fitting capacity.

**Table 2:** The change in energy contribution per residue at the RBD-Bebtelovimab interface and the effect of mutations in BQ.1.1 and XBB.1.5 variants.

| Bebtelovimab-RBD (WT) |  |  |  |  |  | Bebtelovimab-RBD (BQ.1.1) |  |  |  |  |  | Bebtelovimab-RBD (XBB1.5) |  |  |  |  |  |
| --- | --- | --- | --- | --- | --- | --- | --- | --- | --- | --- | --- | --- | --- | --- | --- | --- | --- |
| Type | mAb | RBD | Energy | Dist | BB | Type | mAb | RBD | Energy | Dist | BB | Type | mAb | RBD | Energy | Dist | BB |
| H | Ser30 | Arg346 | -8.5 | 2.87 | b- | IH | Glu53 | Lys440 | -35.07 | 2.71 | -- | H | Ser103 | Lys440 | -0.7 | 3.28 | -- |
| H | Ser32 | Arg346 | -4.1 | 2.83 | -- | H | Arg60 | Val445 | -2.5 | 2.79 | b | IH | Asp56 | Lys444 | -32.43 | 2.84 | -- |
| H | Glu53 | Asn440 | -5.5 | 2.85 | -- | H | Thr96 | Gly446 | -0.5 | 3.45 | bb | IH | Asp58 | Lys444 | -19.7 | 2.73 | -- |
| IH | Asp56 | Lys444 | -27.71 | 2.96 | -- | H | Arg60 | Gly447 | -8.6 | 2.77 | b | H | Arg60 | Pro445 | -6.7 | 2.61 | b |
| IH | Asp58 | Lys444 | -22.05 | 2.77 | -- | H | Asp56 | Asn450 | -8.2 | 2.74 | -- | H | Arg60 | Gly447 | -6.1 | 2.85 | b |
| H | Arg60 | Val445 | -1.6 | 2.84 | b | H | Thr95 | Arg498 | -1.8 | 2.76 | b- | H | Asp56 | Asn450 | -8.6 | 2.81 | -- |
| H | Thr96 | Gly446 | -0.8 | 3.25 | bb | H | Thr96 | Arg498 | -1 | 2.97 | b- | H | Tyr35 | Pro499 | -0.7 | 2.98 | b |
| H | Arg60 | Gly447 | -8.4 | 2.75 | b | H | Asp32 | Thr500 | -4.6 | 2.94 | bb |  |  |  |  |  |  |
| H | Asp56 | Asn450 | -7.2 | 2.7 | -- | H | Asp32 | Gly502 | -0.6 | 3.43 | b |  |  |  |  |  |  |
| H | Thr96 | Gln498 | -4 | 2.81 | b- |  |  |  |  |  |  |  |  |  |  |  |  |
| H | Asp32 | Thr500 | -4.5 | 2.78 | bb |  |  |  |  |  |  |  |  |  |  |  |  |
| H | Gly31 | Thr500 | -0.5 | 3.33 | bb |  |  |  |  |  |  |  |  |  |  |  |  |
| H | Asp32 | Gly502 | -0.5 | 3.44 | b |  |  |  |  |  |  |  |  |  |  |  |  |

## Discussions

Neutralizing antibodies have become an essential form of treatment for individuals who have been diagnosed with severe COVID-19, particularly in cases where vaccination is not a viable option due to high-risk factors. The emergence of new variants has led to a decrease in the efficacy of mAbs in some cases, while in others, mAbs have displayed resistance. This is largely due to the different classifications of mAbs, which are based on the epitopes they recognize on the receptor binding domain of the virus. Some mAbs, such as Sotrovimab, have been observed to retain their neutralization capabilities against certain variants, such as Omicron BA.1, but show a reduced efficacy against others, such as BA.2, BA.4, BA.5, and BA.2.12.1 [2]. Meanwhile, other variants, such as BQ.1.1, have been found to resist the neutralizing effects of mAbs like Sotrovimab, Bebtelovimab, and even combination therapies, such as Bamlanivimab-etesevimab or Evusheld [21]. Additionally, the XBB.1.5 variant, which is considered a leading immune evasion variant, has been estimated to be the most transmissible variant yet [4] and can evade all neutralizing antibodies [21].

The process of mAbs recognition and binding with antigens is highly specific, and even minor changes in the epitope-paratope interface can negatively impact this recognition and binding. With the appearance of new SARS-CoV-2 variants, some mAbs have lost their neutralization ability due to a single amino acid substitution in the epitope, such as the Lys417Asn and Leu452Arg substitutions. However, not all observed epitope mutations result in increased mAb evasion. For example, the R40-1C8 epitope was evaded by the BA.5 variant but not by the BA.1 and BA.2 variants. The current strategies for engineering neutralizing therapeutic mAbs usually involve isolating antibodies from infected or vaccinated individuals or immunized humanized mice [24, 25]. However, with the rapidly evolving SARS-CoV-2 virus, this process has become increasingly challenging, especially where the lead discovery phase must be repeated each time a new variant arises.

Due to the potential of dangerous reinfection associated with new variants of SARS-CoV-2 [3, 26], there is an urgent need to develop broadly active mAbs with both prophylactic and therapeutic potential for high-risk patients [27, 28]. Currently, there is no FDA-approved antibody therapy that effectively neutralizes the latest Omicron variants BQ.1.1 and XBB.1.5. To overcome this challenge, it is possible to use the existing antibody repertoire against all SARS-CoV-2 variants by mapping their epitopes and escape sites. In this study, we used an approach that involves shape complementarity of epitope and paratope residues flexibly, supported by molecular dynamics simulations to confirm stability and resistance to conformational changes. The results suggest that even though BQ.1.1 and XBB.1.5 have multiple new mutations, they are located outside the epitopes of the R200-1F9 and R207-2F11 nAbs. Additionally, the auxiliary contacts made by single amino acid mutations within the epitopes of these mAbs suggest their resilience against both variants. The previously confirmed *in vitro* efficacy of these antibodies further supports the authenticity of the protocol used here [9, 14].

In conclusion, our study suggests that there is potential to redirect existing mAbs against new SARS-CoV-2 variants by mapping their epitopes and escape sites. The approach used in this study, which involves shape complementarity and simulations to confirm stability and resistance to conformational changes, has proven to be effective in identifying mAbs with resilience against both BQ.1.1 and XBB.1.5 variants. Further research is necessary to validate these findings and explore the potential of this approach for the development of broadly active mAbs for the treatment and prevention of COVID-19.

## Methods

### Structures modeling of the Omicron sub-variants and monoclonal antibodies (mAbs)

Among the 126 cross-neutralizing mAbs, Kanika et al. mapped the epitopes of R40-1G8 on the SARS-CoV-2 Spike Wuhan strain and resolved their co-crystal structure (PDB ID: 7SC1) [9]. R40-1G8 was identified as a broadly neutralizing antibody, which was effective against B.1.1.7, B.1.351, B.1.429, B.1.617, and B.1.617.2, as well as 19 prominent potential escape sites in the RBD [9]. However, their recent study demonstrates that this antibody was less or not effective against the Omicron and its sub-variants [14]. To understand the underlying mechanism of this escape, we used the R40-1G8-RBD complex as starting structure and constructed a 3D protein model of R40-1G8-RBD of Omicron BA.1, BA.2, and BA.5 using MOE 2022 package, as described previously [22]. For the construction of isolated RBD structures of Omicron’s sub-variants, the RBD^BA.1^ (PDB ID: 7WBP) crystal structure was used template. Mutations in the Omicron subvariants were made according to the GSAID reported list as displayed in Figure S1A. The structure of an ultra-potent mAb Bebtelovimab bound to Spike protein was retrieved from RSCB PDB (ID: 7MM0) to demonstrate the escape of BQ.1.1 Omicron variant.

The amino acid sequences of all mAbs (variable fragment (FV) regions) including R40-1C8, R207-2F11, and R200-1F9, were retrieved from the coronavirus antibody database (CoV-AbDab) [29] and their CDRs were numbered and annotated in MOE 2022, according to the IMGT system as previously described [30]. The 3D structural model of all mAbs was constructed using default parameters in the MOE antibody modeler package. For each antibody model, the best templates for Framework and CDRs were selected from the built-in antibody database following best scoring, amino acids similarities, and % identity criteria. All constructed models were neutralized in a cubical solvent environment and energy was minimized following the antibody modeling suggested protocol in MOE. Amino acid sequences of the antibodies investigated in this study are available in supplementary data. For Bebtelovimab and related analysis, the crystal structure of mAb-bound RBDWT was retrieved from RSCB PDB (PDB ID: 7MMO) [31].

### Antigen-antibodies docking and epitopes mapping

To map the epitopes of broadly neutralizing mAbs R40-1C8, R207-2F11, and R200-1F9 on SARS-CoV-2 Spike, we used three different protocols to authenticate the outcomes of our analyses. First, a widely implemented DiscoTope (conformational B cell epitope prediction package in IEDB resource) server was used to predict and annotate the potential spatial epitopes on SARS-CoV-2 RBD [32]. Second, the epitope residues suggested by DiscoTope were validated through a linear epitope predictor, Bepipred (IEDB) [33]. Finally, these epitopes were confirmed by corresponding to that reported in actual RBD-mAbs structures of R40-1G8-RBD (ID: 7SC1) and Bebtelovimab-RBD (ID: 7MM0).

To predict the epitopes of R40-1C8, R207-2F11, and R200-1F9, robust antibody docking was used as described previously [20]. For each mAb-RBD complex, 50 conformations were generated by Ag-Ab docking package in Molecular Operating Environment (MOE 2022.02) (Chemical computing group, Montreal, CANADA), where the ligands site were restricted to CDRs and for Ag, RBD was considered as a whole. A protein-ligand interaction fingerprint (PLIF) was generated based on 50 conformations of each mAb which summarized the contribution of each amino acid at the Ag-Ab interface. Based on PLIF results, five epitopes, ranked according to the docking score (kcal/mol), were suggested for each antibody which was further investigated for conservancy in SARS-CoV-2 variants and immunogenicity. As the subvariants neutralization of these mAbs have already been confirmed in vitro [9, 14], mAbs-RBD conformers that were in line with experimental data, bound to conserved epitopes, and establishing significant electrostatic and van der Wall contacts were selected, manually investigated, and further subjected to extensive molecular dynamics simulations for conformational stability and binding affinity estimation.

### Molecular dynamics simulations

Protein models were simulated in a cubic box containing the TIP3P solvent model using GROMACS 2022 under the CHARM36 force field [34]. Models were centered in the box and neutralized with Na and Cl ions, as well as an additional 0.1M concentration of NaCl. The systems were first energy minimized, then equilibrated under constant temperature (NVT) and constant pressure (NPT) conditions for 0.5ns. To prevent systems’ breakage, proteins and solvents were separated and constraints were applied to protein atoms. During the NVT step, the temperature was coupled with the v-rescale (modified Berendsen) thermostat, while the unmodified Berendsen algorithm was used in the NPT step [35]. All systems were simulated for at least 50 ns without structural constraints, using the Particle Mesh Ewald algorithm to calculate long-range electrostatic interactions [36]. After the simulation, artifacts were removed from the MD trajectories using the -PBC and -fit flags in the trjconv tool, along with various functions such as whole, nojump, and rot+trans.

### Free energy perturbation and binding free energies calculation

To calculate binding energies, we used two methods: the endpoint binding free energy MMGBSA method using the HawkDock server [37] and the free energy perturbation method using MMPBSA implemented in GROMACS (versions 5.0 and earlier) [16]. The MMPBSA method is particularly well-suited for calculating the binding energies of various ligands to the same target. The newer versions of GROMACS are not compatible with MMPBSA, so we generated the topology files for each Ab-Ag complex using GROMACS version 5.0. We analyzed an optimized simulation trajectory containing 100 frames to calculate binding free energies, as described in a previous study [20]. Alanine mutagenesis was performed on the DruScorePPI server as described previously [38].

## Declarations

### Authors’ contributions

M.S. and H.W. contributed to the conceptualization of the project. M.S. designed the methodology. M.S. and H.W. wrote the original manuscript draft; H.W. supervised the study and provided funding acquisition.

