## Supplementary Tables and figures for "Assessment of neutralization susceptibility of Omicron subvariants XBB.1.5 and BQ.1.1 against broad-spectrum neutralizing antibodies through epitopes mapping"

### Abstract

The emergence of new variants of the SARS-CoV-2 virus has posed a significant challenge in developing broadly neutralizing antibodies (nAbs) with guaranteed therapeutic potential. Some nAbs, such as Sotrovimab, have exhibited varying levels of efficacy against different variants, while others, such as Bebtelovimab and Bamlanivimab-etesevimab are ineffective against specific variants, including BQ.1.1 and XBB. This highlights the urgent need for developing broadly active mAbs providing prophylactic and therapeutic benefits to high-risk patients, especially in the face of the risk of reinfection from new variants. Here, we aimed to investigate the feasibility of redirecting existing mAbs against new variants of SARS-CoV-2, as well as to understand how BQ.1.1 and XBB.1.5 can evade broadly neutralizing mAbs. By mapping epitopes and escape sites, we discovered that the new variants evade multiple mAbs, including FDA-approved Bebtelovimab, which showed resilience against other Omicron variants. Our approach, which included simulations, free energy perturbations, and shape complementarity analysis, revealed the possibility of identifying mAbs that are effective against both BQ.1.1 and XBB.1.5. We identified two broad-spectrum mAbs, R200-1F9 and R207-2F11, as potential candidates with increased binding affinity to XBB.1.5 and BQ.1.1 compared to the wild-type virus. Additionally, we propose that these mAbs do not interfere with ACE2 and bind to conserved epitopes on the RBD that are not-overlapping, potentially providing a solution to neutralize these new variants either independently or as part of a combination (cocktail) treatment.

### Supplementary Figures

**Figure S1.** (A) Multiple sequence alignment and variations in the SARS-CoV-2 RBD region of Spike protein. (B) Changes in the binding free energy of R40-1C8 bound to the SARS-CoV-2 variants. (C) Changes in the binding free energy of R40-1G8 bound to the SARS-CoV-2 variants.

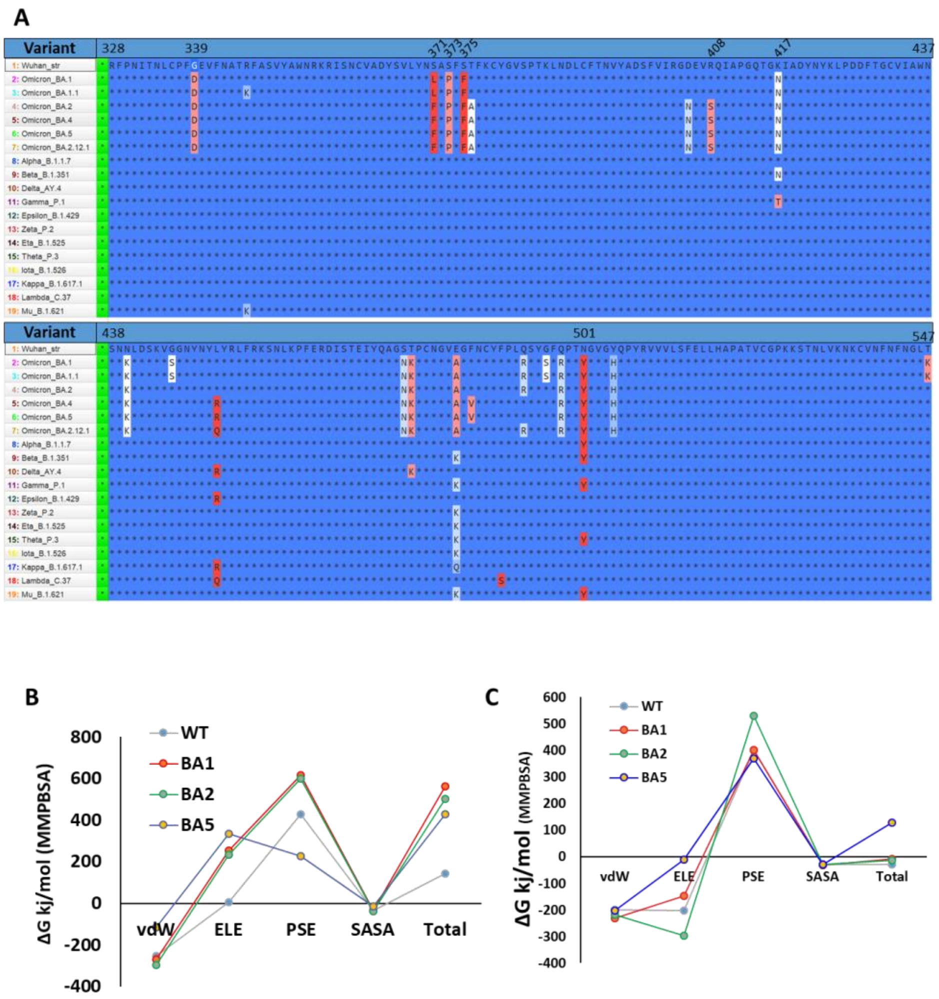

**Figure S2.** The epitope residues on RBD are predicted through DiscoTope (green color cartoon representation) and Molecular Operating Environment (MOE, Cyan color).

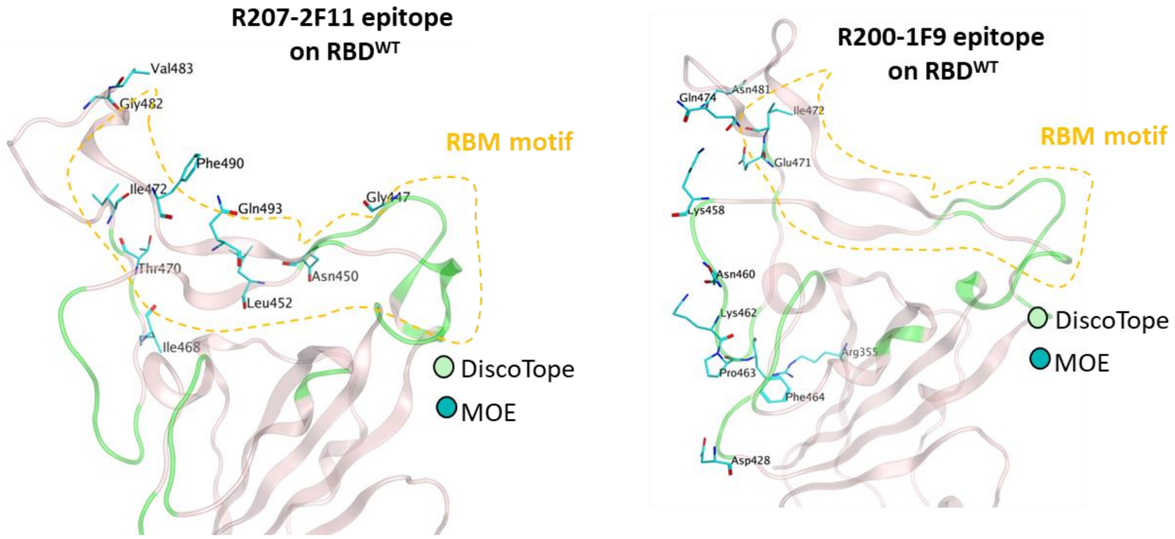

**Figure S3.** (A) Changes in the binding free energy (BFE, *top* MMPBSA, *bottom* MMGBSA) of R200-1F9 bound to the SARS-CoV-2 variants. Plot showing changes in the number of Hydrogen bonds between R200-1F9 and Omicron variants are shown to the right. (B) Changes in the binding free energy (BFE, *top* MMPBSA, *bottom* MMGBSA) of R207-2F11 bound to the SARS-CoV-2 variants. Plot showing changes in the number of Hydrogen bonds between R207-2F11 and Omicron variants are shown to the right.

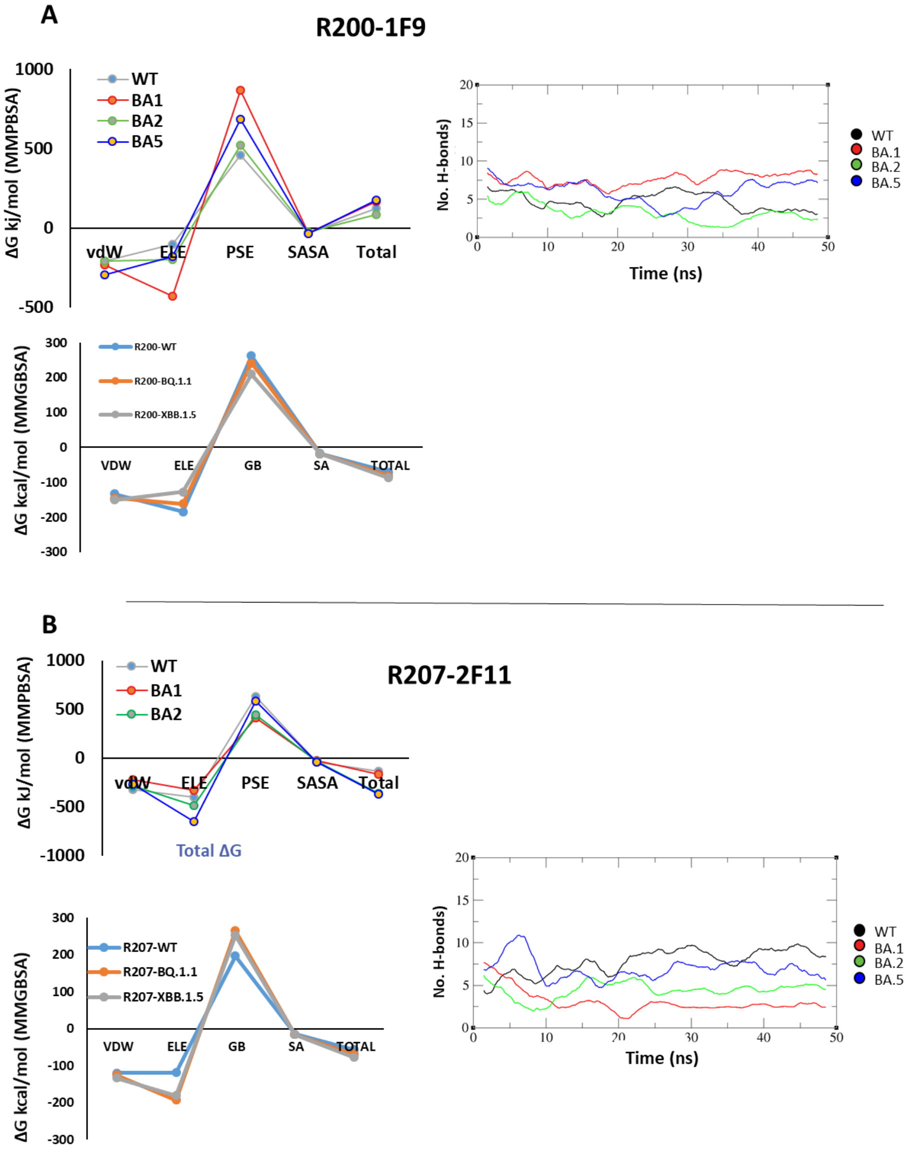

**Figure S4.** Per residues change in binding energy between Bebtelovimab and Omicron BQ.1.1 and XBB.1.5.

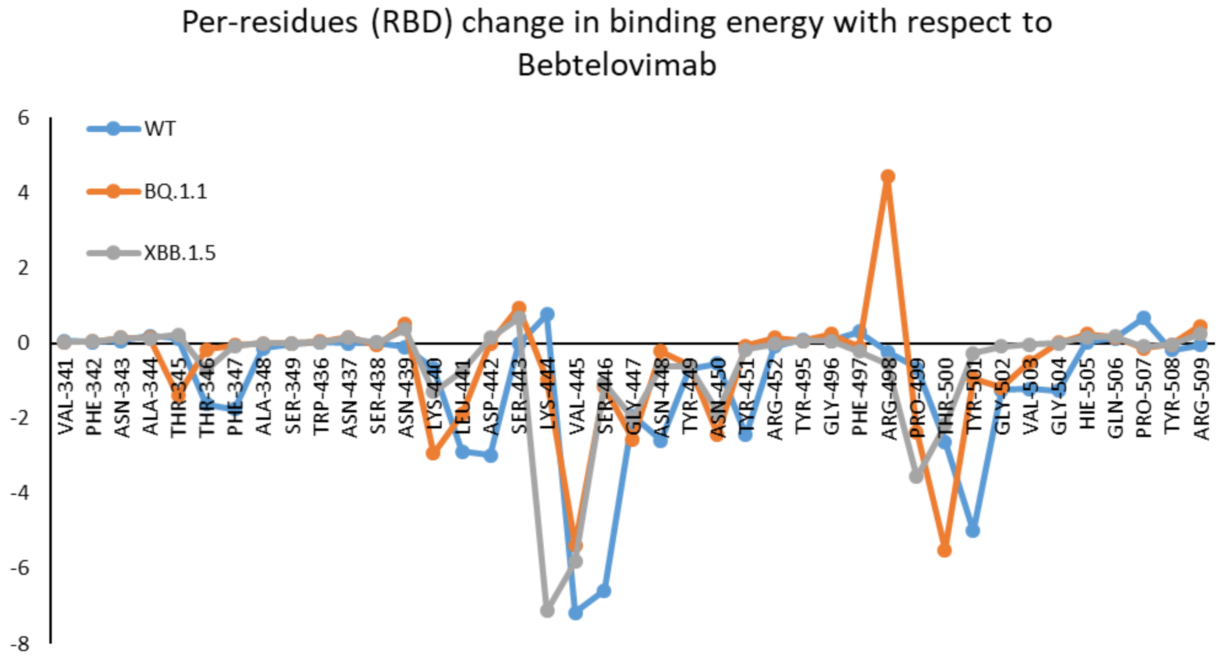

### Supplementary Tables

**Table S1.** The change in energy contribution per residue at the RBD-R401G8 interface and the effect of mutations in BA.1-BA.5 variants.

| R40-1G8-RBD (WT) | | | | | | R40-1G8-RBD (BA1) | | | | | | R40-1G8-RBD (BA2) | | | | | | R40-1G8-RBD (BA5) | | | | | |
|---|---|---|---|---|---|---|---|---|---|---|---|---|---|---|---|---|---|---|---|---|---|---|---|
| Type | mAb | RBD | Energy | Dist | BB | Type | mAb | RBD | Energy | Dist | BB | Type | mAb | RBD | Energy | Dist | BB | Type | mAb | RBD | Energy | Dist | BB |
| H | Ser93 | Arg403 | -2.8 | 3.05 | -- | H | Ser93 | Asp405 | -0.5 | 3.67 | b- | A | Tyr32 | Arg403 | -1.1 | 3.36 | -- | H | Asn92 | Arg403 | -3.5 | 2.87 | b- |
| H | Gln27 | Asp405 | -2.4 | 2.79 | -- | H | Asn92 | Glu406 | -2.6 | 2.85 | -- | H | Gln27 | Asn405 | -3.1 | 2.83 | -- | H | Ser93 | Asn405 | -1.9 | 2.94 | -- |
| IH | Asp94 | Arg408 | -27.7 | 2.73 | -- | IH | Asp94 | Arg408 | -12.93 | 2.7 | -- | H | Asp94 | Asn405 | -3.5 | 2.73 | -- | H | Asp94 | Gln409 | -4 | 2.78 | -- |
| H | Asp94 | Gln409 | -6.4 | 2.81 | -- | H | Asp94 | Gln409 | -6.1 | 2.79 | -- | H | Asn92 | Glu406 | -3.1 | 2.78 | -- | H | Ser56 | Asp420 | -3.1 | 2.67 | -- |
| H | Asn92 | **Lys417** | **-1.2** | **2.97** | **--** | H | Tyr52 | **Asn417** | **-2.4** | **2.69** | -- | H | Asp94 | Gln409 | -4.7 | 2.76 | -- | H | Gly55 | Tyr421 | -1.2 | 2.88 | b- |
| H | Asp94 | **Lys417** | **-7.8** | **2.8** | **b-** | H | Ser56 | Asp420 | -2.4 | 2.63 | -- | H | Tyr52 | Asn417 | -1.8 | 2.78 | -- | H | Asn92 | Tyr453 | -1.5 | 2.89 | -- |
| H | Ser56 | Asp420 | -0.5 | 2.59 | -- | H | Asn92 | Tyr453 | -1.3 | 2.66 | -- | H | Ser56 | Asp420 | -2.9 | 2.77 | -- | A | Tyr33 | Phe456 | -0.5 | 3.81 | -- |
| H | Tyr33 | Leu455 | -1.5 | 2.63 | b | H | Tyr33 | Leu455 | -3.3 | 2.64 | b | H | Asn92 | Tyr453 | -2 | 2.67 | -- | H | Ser31 | Lys458 | -7.4 | 2.77 | -- |
| H | Ser31 | Lys458 | -7.5 | 2.78 | -- | H | Ser30 | Lys458 | -0.5 | 2.79 | b- | H | Tyr33 | Leu455 | -3.4 | 2.67 | b | H | Ser31 | Tyr473 | -0.9 | 2.82 | b- |
| H | Ser56 | Asn460 | -2.4 | 2.84 | -- | H | Ser31 | Lys458 | -6.1 | 2.81 | -- | H | Ser31 | Lys458 | -5 | 2.79 | -- | H | Asn32 | Ala475 | -4.9 | 2.79 | b |
| H | Ser31 | Gln474 | -2.9 | 2.99 | b- | H | Ser31 | Tyr473 | -3.2 | 2.73 | b- | H | Gly55 | Asn460 | -2.1 | 2.9 | b- | H | Gly26 | Asn477 | -5.1 | 2.99 | b* |
| H | Arg97 | Phe486 | -8.3 | 2.86 | b | H | Ser31 | Gln474 | -3.1 | 2.65 | b | H | Ser31 | Tyr473 | -3.1 | 2.75 | b- | H | Tyr32 | Tyr495 | -2 | 2.71 | b |
| H | Gly26 | Asn487 | -3.3 | 2.69 | b- | H | Asn32 | Ala475 | -1.7 | 2.8 | b | A | Val2 | Phe486 | -0.6 | 4.5 | -- | H | Ser31 | Tyr501 | -1.7 | 2.81 | -- |
| H | Ser30 | Asn501 | -2.1 | 2.84 | -- | H | Gly26 | Asn477 | -2.5 | 3.07 | bb | H | Ser31 | Asn487 | -2.1 | 2.71 | -- | H | Gly28 | His505 | -4.1 | 2.82 | b- |
| H | Asn92 | Tyr505 | -2.1 | 2.7 | b- | H | Leu27 | Asn477 | -1.1 | 2.86 | b- | H | Asp98 | Tyr489 | -5 | 2.63 | -- | | | | | | |
| | | | | | | H | Tyr32 | Ser494 | -3.9 | 2.71 | b | A | Phe102 | Arg493 | -0.5 | 3.43 | -- | | | | | | |
| | | | | | | | | | | | | H | Tyr32 | Ser494 | -2.4 | 2.67 | b | | | | | | |
| | | | | | | | | | | | | H | Ser30 | Tyr495 | -0.5 | 3.12 | b | | | | | | |
| | | | | | | | | | | | | H | Ser31 | Arg498 | -3 | 3 | -- | | | | | | |
| | | | | | | | | | | | | H | Gly28 | Gly502 | -2.4 | 3.03 | bb | | | | | | |

**Table S2.** The change in energy contribution per residue at the RBD-R401C8 interface and the effect of mutations in all Omicron variants.

| R40-1C8-RBD (WT) | | | | | | R40-1C8-RBD (BA1) | | | | | |
|---|---|---|---|---|---|---|---|---|---|---|---|
| Type | mAb | RBD | Energy | Dist | BB | Type | mAb | RBD | Energy | Dist | BB |
| H | Tyr102 | Glu406 | -4.9 | 2.6 | -- | H | Thr53 | Arg403 | -0.6 | 3.24 | -- |
| H | Tyr33 | Gly413 | -2.8 | 2.93 | b- | H | Tyr102 | Glu406 | -4.1 | 2.78 | -- |
| A | Tyr49 | Lys417 | -0.7 | 4.6 | -- | H | Leu99 | Arg408 | -2.5 | 3.19 | b- |
| IH | Asp105 | Lys417 | -25.01 | 2.7 | -- | H | Tyr33 | Gly413 | -3.4 | 2.75 | b- |
| IH | Arg97 | Asp420 | -15.25 | 3.07 | -- | H | Asp98 | Thr415 | -1.3 | 2.95 | b- |
| H | Glu1 | Lys458 | -3 | 3.06 | b- | IH | Arg97 | Asp420 | -22.09 | 2.85 | -- |
| H | Asn32 | Asn460 | -6.7 | 2.86 | -- | H | Glu1 | Lys458 | -9.7 | 2.77 | b- |
| A | Ser60 | Phe486 | -0.8 | 4.64 | b- | H | Asn32 | Asn460 | -6.5 | 2.86 | -- |
| H | Val58 | Tyr489 | -2.1 | 2.79 | b- | A | Ser60 | Phe486 | -0.7 | 3.94 | b- |
| H | Thr53 | Tyr505 | -2.3 | 2.65 | -- | H | Val58 | Tyr489 | -4 | 2.7 | b- |
| | | | | | | H | Gly50 | His505 | -1.8 | 2.73 | b- |

| R40-1C8-RBD (BA2) | | | | | | R40-1C8-RBD (BA5) | | | | | |
|---|---|---|---|---|---|---|---|---|---|---|---|
| Type | mAb | RBD | Energy | Dist | BB | Type | mAb | RBD | Energy | Dist | BB |
| H | Thr53 | Arg403 | -2.6 | 2.89 | -- | H | Thr53 | Arg403 | -3.8 | 2.88 | -- |
| H | Asp101 | Asn405 | -1.8 | 2.76 | -- | H | Asp101 | Asn405 | -3.4 | 2.74 | -- |
| H | Tyr102 | Glu406 | -1.2 | 2.54 | -- | A | Tyr102 | Asn405 | -0.7 | 3.6 | -- |
| H | Tyr33 | Gly413 | -2.6 | 2.66 | b- | H | Tyr102 | Glu406 | -4.3 | 2.62 | -- |
| H | Arg97 | Thr415 | -2 | 2.98 | -- | H | Tyr33 | Gly413 | -3.1 | 2.77 | b- |
| IH | Arg97 | Asp420 | -11.71 | 3.27 | -- | H | Tyr33 | Thr415 | -0.9 | 3 | b- |
| H | Glu1 | Lys458 | -9 | 2.91 | b- | H | Arg97 | Thr415 | -2.6 | 2.87 | -- |
| H | Asn32 | Asn460 | -6.6 | 2.86 | -- | IH | Arg97 | Asp420 | -11.66 | 2.7 | -- |
| A | Ser60 | Phe486 | -1.7 | 4.37 | b- | H | Glu1 | Lys458 | -8.6 | 2.79 | b- |
| H | Val58 | Tyr489 | -4.5 | 2.69 | b- | H | Asn32 | Asn460 | -6.1 | 2.92 | -- |
| H | Asp101 | His505 | -10 | 2.91 | -- | H | Val58 | Tyr489 | -3.5 | 2.67 | b- |
| | | | | | | H | Asp101 | His505 | -8 | 2.82 | -- |

| R40-1C8-RBD (BQ.1.1) | | | | | | R40-1C8-RBD (XBB.1.5) | | | | | |
|---|---|---|---|---|---|---|---|---|---|---|---|
| Type | mAb | RBD | Energy | Dist | BB | Type | mAb | RBD | Energy | Dist | BB |
| H | Thr53 | Arg403 | -3.2 | 2.97 | -- | H | Thr53 | Arg403 | -3.2 | 2.97 | -- |
| H | Asp101 | Asn405 | -2.5 | 2.75 | -- | H | Asp101 | Asn405 | -2.5 | 2.75 | -- |
| A | Tyr102 | Asn405 | -0.7 | 3.86 | -- | A | Tyr102 | Asn405 | -0.7 | 3.86 | -- |
| H | Tyr102 | Glu406 | -4 | 2.61 | -- | H | Tyr102 | Glu406 | -4 | 2.61 | -- |
| H | Tyr33 | Gly413 | -3.2 | 2.7 | b- | H | Tyr33 | Gly413 | -3.2 | 2.7 | b- |
| H | Tyr33 | Thr415 | -1 | 3.29 | b- | H | Tyr33 | Thr415 | -1 | 3.29 | b- |
| H | Arg97 | Thr415 | -3 | 2.9 | -- | H | Arg97 | Thr415 | -3 | 2.9 | -- |
| IH | Arg97 | Asp420 | -12.44 | 2.79 | -- | IH | Arg97 | Asp420 | -12.44 | 2.79 | -- |
| H | Glu1 | Lys458 | -5.8 | 3 | b- | H | Glu1 | Lys458 | -5.8 | 3 | b- |
| H | Val58 | Tyr489 | -4.1 | 2.66 | b- | H | Val58 | Tyr489 | -4.1 | 2.66 | b- |
| H | Asp101 | His505 | -8.2 | 2.82 | -- | H | Asp101 | His505 | -8.2 | 2.82 | -- |

**Table S3.** The change in energy contribution per residue at the RBD-R200-1F9 interface and the effect of mutations in all Omicron variants.

| R200-1F9-RBD (WT) | | | | | | R200-1F9-RBD (BA1) | | | | | |
|---|---|---|---|---|---|---|---|---|---|---|---|
| Type | mAb | RBD | Energy | Dist | BB | Type | mAb | RBD | Energy | Dist | BB |
| H | Asn1 | Arg355 | -3.5 | 3.11 | *- | H | Gln27 | Arg355 | -9.1 | 2.79 | -- |
| H | Gln27 | Arg355 | -6.7 | 2.88 | -- | H | Ser28 | Asp428 | -4.4 | 2.82 | -- |
| H | Ser28 | Asp428 | -3.3 | 2.61 | -- | H | Ser57 | Lys458 | -2.4 | 2.96 | b |
| H | Arg101 | Lys458 | -6 | 2.87 | b- | H | Arg101 | Lys458 | -4.1 | 2.74 | b |
| H | His104 | Asn460 | -3.6 | 3.01 | b- | H | His104 | Asn460 | -1 | 3.01 | b |
| H | Ser105 | Lys462 | -0.6 | 3.15 | -- | H | His104 | Lys462 | -6.4 | 2.94 | b- |
| H | Asn93 | Pro463 | -3.8 | 2.91 | b- | H | Ser105 | Lys462 | -0.6 | 3.25 | -- |
| H | Asn93 | Phe464 | -1 | 2.77 | b- | H | Asn1 | Phe464 | -1 | 2.79 | bb |
| IH | Arg58 | Glu471 | -30.8 | 2.97 | -- | IH | Asp62 | Arg466 | -14.1 | 3.13 | -- |
| H | Tyr60 | Glu471 | -6 | 2.66 | -- | H | Trp94 | Arg466 | -2.8 | 3.16 | bb |
| H | Ser56 | Ile472 | -2.3 | 2.71 | b- | IH | Arg58 | Glu471 | -32.9 | 2.91 | -- |
| H | Ser56 | Gln474 | -1.3 | 3.15 | b- | H | Tyr60 | Glu471 | -4.7 | 2.67 | -- |
| H | Gly55 | Asn481 | -1.4 | 3 | b- | H | Ser56 | Ile472 | -2.4 | 2.66 | b |
| H | Arg58 | Asn481 | -5.8 | 2.74 | -- | H | Ser56 | Gln474 | -2.2 | 2.9 | b |
| | | | | | | H | Gly55 | Asn481 | -1.7 | 2.8 | b- |

| R200-1F9-RBD (BA2) | | | | | | R200-1F9-RBD (BA5) | | | | | |
|---|---|---|---|---|---|---|---|---|---|---|---|
| Type | mAb | RBD | Energy | Dist | BB | Type | mAb | RBD | Energy | Dist | BB |
| H | Gln27 | Arg355 | -9.1 | 2.79 | -- | H | Gln27 | Arg355 | -9.1 | 2.79 | -- |
| H | Ser28 | Asp428 | -4.5 | 2.82 | -- | H | Ser28 | Asp428 | -4.4 | 2.82 | -- |
| H | Ser57 | Lys458 | -2.4 | 2.96 | b | H | Ser57 | Lys458 | -2.4 | 2.96 | b |
| H | Arg101 | Lys458 | -4.1 | 2.74 | b | H | Arg101 | Lys458 | -4.1 | 2.74 | b |
| H | His104 | Asn460 | -1 | 3.02 | b | H | His104 | Asn460 | -1 | 3.01 | b |
| H | His104 | Lys462 | -6.4 | 2.94 | b- | H | His104 | Lys462 | -6.4 | 2.94 | b- |
| H | Ser105 | Lys462 | -0.6 | 3.25 | -- | H | Ser105 | Lys462 | -0.6 | 3.25 | -- |
| H | Asn1 | Phe464 | -0.9 | 2.79 | bb | H | Asn1 | Phe464 | -1 | 2.79 | bb |
| IH | Asp62 | Arg466 | -14.1 | 3.13 | -- | IH | Asp62 | Arg466 | -14.2 | 3.12 | -- |
| H | Trp94 | Arg466 | -2.8 | 3.16 | bb | H | Trp94 | Arg466 | -2.8 | 3.16 | bb |
| IH | Arg58 | Glu471 | -32.9 | 2.91 | -- | IH | Arg58 | Glu471 | -33 | 2.91 | -- |
| H | Tyr60 | Glu471 | -4.7 | 2.67 | -- | H | Tyr60 | Glu471 | -4.7 | 2.67 | -- |
| H | Ser56 | Ile472 | -2.4 | 2.66 | b | H | Ser56 | Ile472 | -2.4 | 2.66 | b |
| H | Ser56 | Gln474 | -2.2 | 2.91 | b | H | Ser56 | Gln474 | -2.2 | 2.91 | b |
| H | Gly55 | Asn481 | -1.7 | 2.8 | b- | H | Gly55 | Asn481 | -1.7 | 2.8 | b- |

| R200-1F9-RBD (BQ.1.1) | | | | | | R200-1F9-RBD (XBB.1.5) | | | | | |
|---|---|---|---|---|---|---|---|---|---|---|---|
| Type | mAb | RBD | Energy | Dist | BB | Type | mAb | RBD | Energy | Dist | BB |
| H | Gln27 | Arg355 | -9.1 | 2.78 | -- | H | Asn1 | Phe464 | -1.3 | 2.83 | bb |
| H | Ser28 | Asp428 | -2.1 | 2.53 | -- | H | Gln27 | Arg355 | -9 | 2.82 | -- |
| H | Ser30 | Asp428 | -4.6 | 2.63 | -- | H | Ser28 | Asp428 | -1.7 | 2.56 | -- |
| H | Ser57 | Lys458 | -2.4 | 2.96 | b | H | Gly55 | Asn481 | -1.2 | 2.73 | b- |
| H | Arg101 | Lys458 | -2.8 | 2.7 | b | H | Ser56 | Ile472 | -2.3 | 2.65 | b |
| H | His104 | Lys460 | -3.9 | 3.12 | b | H | Ser56 | Gln474 | -2.1 | 2.88 | b |
| H | His104 | Lys462 | -1.1 | 3.57 | b- | H | Ser56 | Asn481 | -2.4 | 3.14 | b* |
| IH | Asp62 | Arg466 | -19.5 | 2.8 | -- | H | Ser57 | Lys458 | -2 | 2.99 | b |
| H | Trp94 | Arg466 | -1.6 | 3.25 | bb | IH | Arg58 | Glu471 | -30.5 | 2.95 | -- |
| IH | Arg58 | Glu471 | -33.2 | 2.87 | -- | H | Tyr60 | Glu471 | -6.2 | 2.69 | -- |
| H | Tyr60 | Glu471 | -5.3 | 2.67 | -- | IH | Asp62 | Arg466 | -15.3 | 2.71 | -- |
| H | Ser56 | Ile472 | -2.2 | 2.76 | b | H | Ser71 | Asn481 | -0.7 | 3.4 | b- |
| H | Ser56 | Gln474 | -2.7 | 2.94 | b | H | Asn93 | Pro463 | -0.8 | 3.11 | b |
| H | Gly55 | Asn481 | -2.9 | 2.76 | b- | H | Trp94 | Arg466 | -2.7 | 3.2 | bb |
| | | | | | | H | Arg101 | Lys458 | -3.2 | 2.77 | b |
| | | | | | | H | His104 | Lys460 | -4.2 | 3.13 | b |
| | | | | | | H | His104 | Lys462 | -8.8 | 2.79 | b- |

**Table S4.** The change in energy contribution per residue at the RBD-R207-2F11 interface and the effect of mutations in all Omicron variants.

R207-2F11-RBD (WT)

| Type | mAb | RBD | Energy | Dist | BB |
|---|---|---|---|---|---|
| A | Tyr101 | Ser349 | -0.6 | 3.78 | b |
| H | Lys31 | Gly447 | -9.4 | 2.73 | b |
| H | Tyr49 | Asn450 | -3.7 | 2.97 | b- |
| H | Asp50 | Asn450 | -1 | 3.09 | b |
| H | Tyr91 | Asn450 | -0.6 | 3.38 | b |
| H | Tyr101 | Asn450 | -4.3 | 2.83 | b- |
| H | Tyr52 | Ile468 | -1.8 | 2.81 | b |
| H | Asn93 | Ile472 | -3.4 | 2.94 | b |
| H | Lys1 | Gly482 | -10.3 | 2.72 | bb |
| H | Asn30 | Gln493 | -2.1 | 2.92 | -- |
| H | Tyr32 | Gln493 | -2.1 | 2.57 | -- |

R207-2F11-RBD (BA1)

| Type | mAb | RBD | Energy | Dist | BB |
|---|---|---|---|---|---|
| H | Tyr49 | Arg346 | -0.7 | 2.94 | -- |
| A | Tyr101 | Ser349 | -0.5 | 3.61 | b |
| H | Ser52 | Ser446 | -1.2 | 2.89 | -- |
| H | Lys31 | Gly447 | -10.9 | 2.73 | b |
| H | Tyr49 | Asn450 | -3.7 | 2.93 | b- |
| H | Tyr91 | Asn450 | -1.3 | 3.24 | b |
| H | Tyr101 | Asn450 | -5.1 | 2.85 | b- |
| H | Asn93 | Ile472 | -2.6 | 2.98 | b |
| H | Lys1 | Asn481 | -2 | 3.02 | bb |
| H | Lys1 | Gly482 | -8.3 | 2.73 | bb |
| H | Asn30 | Arg493 | -5.6 | 2.76 | -- |
| H | Tyr32 | Ser494 | -1.1 | 2.79 | -- |

R207-2F11-RBD (BA2)

| Type | mAb | RBD | Energy | Dist | BB |
|---|---|---|---|---|---|
| H | Tyr101 | Ala352 | -1.4 | 3.15 | b |
| H | Lys31 | Gly447 | -9.3 | 2.71 | b |
| H | Asn53 | Asn448 | -0.5 | 2.96 | -- |
| A | Lys31 | Tyr449 | -0.5 | 4.61 | -- |
| H | Tyr49 | Asn450 | -4.3 | 2.9 | b- |
| H | Tyr101 | Asn450 | -3 | 2.87 | b- |
| H | Asn93 | Ile472 | -0.8 | 3.23 | b |
| H | Leu94 | Asn481 | -3.2 | 2.9 | b- |
| H | Lys1 | Gly482 | -7.1 | 2.75 | bb |
| H | Asn30 | Arg493 | -5.8 | 2.83 | -- |
| H | Tyr32 | Ser494 | -1.1 | 2.78 | -- |

R207-2F11-RBD (BA5)

| Type | mAb | RBD | Energy | Dist | BB |
|---|---|---|---|---|---|
| H | Tyr101 | Ala352 | -1.4 | 3.16 | b |
| H | Lys31 | Gly447 | -8.3 | 2.72 | b |
| H | Tyr49 | Asn450 | -4.4 | 2.91 | b- |
| H | Tyr101 | Asn450 | -2.9 | 2.87 | b- |
| H | Tyr91 | Arg452 | -4.1 | 2.67 | b- |
| IH | Asp92 | Arg452 | -5.69 | 3.36 | *- |
| H | Asn93 | Ile472 | -0.5 | 3.28 | b |
| H | Leu94 | Asn481 | -3 | 2.92 | b- |
| H | Lys1 | Gly482 | -6.2 | 2.78 | bb |
| H | Asn30 | Gln493 | -2 | 2.88 | -- |
| H | Tyr32 | Gln493 | -1.7 | 2.59 | -- |
| H | Tyr32 | Ser494 | -1.1 | 2.73 | -- |

R207-2F11-RBD (BA.1.1)

| Type | mAb | RBD | Energy | Dist | BB |
|---|---|---|---|---|---|
| H | Tyr101 | Ala352 | -2 | 2.95 | b |
| H | Lys31 | Gly447 | -7.7 | 2.72 | b |
| H | Asn53 | Asn448 | -1 | 2.93 | -- |
| H | Asp50 | Tyr449 | -2.6 | 2.89 | b |
| H | Tyr49 | Asn450 | -1.8 | 3.26 | b- |
| H | Tyr101 | Asn450 | -3.8 | 2.84 | b- |
| H | Tyr91 | Arg452 | -3.4 | 2.7 | b- |
| H | Asp92 | Arg452 | -6.5 | 2.86 | b- |
| H | Tyr52 | Ile468 | -2.5 | 2.86 | b |
| A | Asn93 | Tyr473 | -0.9 | 3.78 | -- |
| H | Lys1 | Asn481 | -8.4 | 2.98 | bb |
| H | Asn30 | Gln493 | -2.1 | 2.87 | -- |
| H | Tyr32 | Gln493 | -1.3 | 2.62 | -- |
| H | Tyr32 | Ser494 | -1.1 | 2.74 | -- |

R207-2F11-RBD (XBB.1.5 )

| Type | mAb | RBD | Energy | Dist | BB |
|---|---|---|---|---|---|
| H | Tyr101 | Ala352 | -0.8 | 3.33 | b |
| H | Lys31 | Gly447 | -6.8 | 2.71 | b |
| H | Asn53 | Gly447 | -1.5 | 3.06 | b |
| H | Asp50 | Tyr449 | -1.3 | 3.11 | b |
| H | Asp50 | Asn450 | -2.4 | 3.09 | b |
| H | Tyr101 | Asn450 | -3.3 | 2.76 | b- |
| A | Tyr32 | Leu452 | -0.6 | 3.99 | -- |
| H | Tyr52 | Ile468 | -2.2 | 2.85 | b |
| H | Asn93 | Ile472 | -1.5 | 2.8 | b |
| A | Asn93 | Tyr473 | -0.7 | 3.65 | -- |
| H | Lys1 | Gly482 | -10.4 | 2.93 | bb |
| H | Gln27 | Val483 | -0.6 | 3.42 | b |
| H | Asp92 | Ser490 | -2.5 | 2.83 | -- |
| H | Asn93 | Ser490 | -2.5 | 2.98 | -- |
| H | Asn30 | Gln493 | -1.9 | 2.9 | -- |
| H | Tyr32 | Gln493 | -1.4 | 2.56 | -- |
| H | Tyr32 | Ser494 | -0.9 | 2.8 | -- |
